# Transkingdom Analysis of the Female Reproductive Tract Reveals Bacteriophages Form Communities

**DOI:** 10.1101/2021.10.12.464088

**Authors:** Ferralita S. Madere, Michael Sohn, Angelina Winbush, Breóna Barr, Alex Grier, James Java, Tracy Meiring, Anna-Lise Williamson, Linda-Gail Bekker, David H. Adler, Cynthia L. Monaco

**Affiliations:** Department of Microbiology and Immunology, University of Rochester Medical Center, Rochester, NY, USA; Department of Biostatistics and Computational Biology, University of Rochester Medical Center, Rochester, NY, USA; University of Rochester School of Medicine & Dentistry, Rochester, NY, USA; Department of Rural Family Medicine, West Virginia University, Morgantown, WV; UR Genomics Research Center, University of Rochester Medical Center, Rochester, NY, USA; Institute of Infectious Diseases & Molecular Medicine and Division of Medical Virology, Faculty of Health Sciences, University of Cape Town, Anzio Road, Observatory, Cape Town, South Africa; National health Laboratory Service, Groote Schuur Hospital, Cape Town, South African; Desmond Tutu HIV Centre, Institute of Infectious Diseases & Molecular Medicine, Faculty of Health Sciences, University of Cape Town, Anzio Road, Observatory, Cape Town, South Africa; Department of Emergency Medicine, University of Rochester Medical Center, Rochester, NY, USA; Department of Internal Medicine, Division of Infectious Diseases, University of Rochester Medical Center, Rochester, NY, USA

**Keywords:** virome, microbiome, bacterial vaginosis, bacteriophage, transkingdom associations, female genital tract

## Abstract

The female reproductive tract (FRT) microbiome plays an important role in maintaining vaginal health. Viruses play a key role in regulating other microbial ecosystems, but little is known about how the FRT viruses (virome), particularly bacteriophages, impacts FRT health and dysbiosis. We hypothesize that bacterial vaginosis is associated with alterations in the FRT virome, and these changes correlate with bacteriome shifts. We conducted a retrospective, longitudinal analysis of vaginal swabs collected from 54 bacterial vaginosis (BV)-positive and 46 BV-negative South African women. Bacteriome analysis revealed samples clustered into five distinct bacterial community groups (CG). Bacterial alpha diversity was significantly associated with BV. Virome analysis on a subset of baseline samples showed FRT bacteriophages clustering into novel viral state types (VSTs), a viral community clustering system based on virome composition and abundance. Distinct BV bacteriophage signatures included increased alpha diversity along with *Bacillus, Burkholderia* and *Escherichia* bacteriophages. Discriminate bacteriophage-bacteria transkingdom associations were also identified between *Bacillus* and *Burkholderia* viruses and BV-associated bacteria, providing key insight for future studies elucidating transkingdom interactions driving BV-associated microbiome perturbations. In this cohort, bacteriophage-bacterial associations suggest complex interactions which may play a role in the establishment and maintenance of BV.

## Introduction

The female reproductive tract (FRT) houses a compositionally dynamic environment where the host participates in an intricate interplay with a microbiome composed of bacteria and archaea (bacteriome), fungi (fungome), viruses (virome), and occasional prokaryotic parasites [1, 2]. The FRT microbiome plays an important protective role in maintaining vaginal health and preventing urogenital diseases such as bacterial vaginosis (BV), yeast infections, pre-term birth and sexually transmitted infections including HIV [3-5]. Prior studies of the FRT microbiome have primarily focused on determining bacterial composition and function. At least five different bacterial community groupings have been described within the FRT, distinguishable by the dominance of *Lactobacillus* species or presence of more diverse anaerobes [6, 7]. Prevalence of these communities varies by ethnic group, with majority of Caucasian women hosting *Lactobacillus*-dominant FRT microbiomes, whereas African women tend to be asymptomatically colonized by higher diversity FRT microbiota [7, 8]. *Lactobacillus-*dominant FRT bacteriomes, especially *L. crispatus*, protect against vaginal diseases by competitive exclusion against pathogenic bacteria for space and nutrients or by promoting an acidic vaginal environment via production of lactic acid and maintaining a low inflammatory state [9-11]. BV, the most common cause of vaginal discharge in reproductive-age women, is a symptomatic clinical condition characterized by a shift in the FRT microbiota away from a low inflammatory, *Lactobacillus*-dominant microbiome to more diverse community including facultative anaerobes. BV is associated with an increased risk of sexually transmitted infection (STI) acquisition and pre-term birth [12, 13]. FRT bacteriome shifts can occur rapidly and may be related to shifts in bacteriophage populations [14, 15]. Specific BV-associated bacteria include *Gardnerella vaginalis, Prevotella, Fusobacterium, Atopobium vaginae, Megasphaera*, and *Sneathia* among others [16].

Viruses rival bacterial numbers in the microbiome and are more diverse [1]. However, studies of the virome have been limited in part due to lack of a common viral genetic element analogous to the bacterial 16S rRNA gene, as well as the high genetic diversity between viral species [1, 17]. The FRT virome is home to eukaryotic viruses and bacteriophages [1]. Compared to the FRT bacteriome, little is known about the viral communities of the FRT and how their interactions with bacteria contribute to disease states such as bacterial vaginosis (BV). The few prior studies that have examined the FRT virome have mainly concentrated on the DNA eukaryotic virome, finding *Papillomaviridae, Polyomaviridae, Herpesviridae, Poxviridae, Adenoviridae*, and *Anelloviridae* present [18, 19]. However, bacteriophages are the largest and most abundant viral group and can modulate bacterial composition and abundance, suggesting that they may play an important role in regulating bacterial composition of the FRT microbiome [17, 20]. Bacteriophages may be lytic, hijacking bacterial host replication machinery in order to replicate and then lysing the bacterial host to release virions, or lysogenic, whereby bacteriophage DNA is integrated into the bacterial host genome as a prophage and replicates with bacterial genome replication [21]. This lysogenic lifestyle can result in generalized transduction of bacterial genes between bacterial hosts that can confer increased fitness through methods such as toxin production, carbohydrate metabolism, or antibiotic resistance [22]. Upon environmental stress, lysogenic bacteria can become lytic, and therefore may serve to regulate bacterial populations in unfavorable host conditions.

Data on bacteriophage populations in the FRT are limited. One study revealed numerous *Caudovirales* order bacteriophage in the FRT; however, no relationship was found between FRT *Caudovirales* bacteriophages and bacterial populations [4]. Additionally, other investigations studying BV-positive and BV-negative samples in a cohort of Danish women found no significant difference in viral nor bacterial alpha diversity between BV-positive and BV-negative women [23]. While brief characterization of the vaginal microbiome in pregnant women found an absence of *Microviridae* and *Herelleviridae* bacteriophage families in a pregnant woman without vaginitis [24]. Neither of these studies, however, employed significant sequencing depths necessary to perform a through characterization of the FRT phageome and identify distinct perturbations between health and disease. *Lactobacillus* bacteriophages have been isolated from human FRT *Lactobacillus* species and have been observed to have broad host ranges, capable of infecting multiple *Lactobacillus* species [24]. Upon induction, lysogenic phages within lactobacilli can lyse their host and enable pathogenic bacteria to flourish, suggesting their possible role in regulating bacteriome composition and promoting the growth of BV-associated bacteria [25-28]. Further in support of this hypothesis, high gene expression of the CRISPR anti-bacteriophage defense system occurs in BV, suggesting that an altered phage load could contribute to the hallmark dysbiosis observed in BV [29-32].

Herein we investigated FRT virome composition and transkingdom bacterial-bacteriophage associations within the FRT. We assessed the vaginal samples of 100 young, sexually active BV-positive and -negative South African women to identify discriminate viral and bacterial signatures in health and FRT disease. We show for the first time that FRT DNA bacteriophage populations cluster into community groupings that correspond to bacterial community groupings, and that specific bacteriophages correlate with bacteria associated with and protective against BV. These findings improve our understanding of the transkingdom associations in the FRT microbiome and the impact that these could have on the induction and pathogenesis of disease.

## Methods

### Study Cohort

De-identified vaginal swabs from the University of Cape Town HPV-HIV study [33] were retrospectively used for bacteriome and virome analysis. This cohort was comprised of 50 HIV-positive and 50 HIV-negative young, sexually active women between the ages of 16 and 21 recruited from the youth community center and clinic in two urban disadvantaged communities (Masiphumelele and Mthatha townships) in Cape Town, South Africa between October 2012 and October 2014. Informed consent was obtained from all participants above 18 years of age and parental consent was obtained for participants of age 17 or younger. This study was approved by the Institutional Review Board at University of Rochester Medical Center and the Human Research Ethics committee at the University of Cape Town. Vaginal swabs were self-collected approximately every 6 months by subjects using Dacron swabs high within the vagina, placed in Digene transport media, and frozen at −80°C until use. Pap smears were taken at baseline and at least one other visit, and tests for HPV, *Trichomonas*, and BV performed [33]. HIV status was confirmed upon enrollment, and CD4+ T cell count for those who were HIV-positive was determined. Exclusion criteria included prior HPV vaccination or cervical surgery. Sexual history, recent contraceptive methods and current HIV treatment interventions were also queried at study entry.

### Bacterial 16S rRNA gene amplicon sequencing

Total nucleic acid was extracted from 253 resuspended vaginal swab samples using the MagNA Pure Compact Nucleic Acid Isolation Kit (Roche) [33]. 16S rRNA gene amplicon sequencing was performed with primers specific to the V3-V4 region [34] followed by amplicon pooling, bead-based normalization and sequencing on the Illumina MiSeq platform at 312 bp paired-end reads (University of Rochester Genomics Research Center, UR GRC). Water processed similarly to samples and pre-defined bacterial mixtures (Zymo) were run as negative and positive controls, respectively. Eleven samples failed 16S rRNA gene amplification.

### Bacterial 16S rRNA gene amplicon analysis

Raw data from the Illumina MiSeq was first converted into FASTQ format 2 × 312 paired-end sequence files using the bcl2fastq program (v1.8.4) provided by Illumina. Format conversion was performed without de-multiplexing, and the EAMMS algorithm was disabled. All other settings were default. Reads were multiplexed using a configuration described previously [34]. The extract_barcodes.py script from QIIME (v1.9.1) [35] was used to split read and barcode sequences into separate files suitable for import into QIIME 2 (v2018.11) [36], which was used to perform all subsequent read processing and characterization of sample composition. Reads were demultiplexed requiring exact barcode matches, and 16S primers were removed allowing 20% mismatches and requiring at least 18 bases. Cleaning, joining, and denoising were performed using DADA2 [37]: reads were truncated (forward reads to 260 bps and reverse reads to 240 bps), error profiles were learned with a sample of one million reads per run, and a maximum expected error of two was allowed. Taxonomic classification was performed with custom naïve Bayesian classifiers trained on target-region specific subsets of the August, 2013 release of GreenGenes [38]. Sequence variants that could not be classified to at least the phylum level were discarded. Sequencing variants observed fewer than ten times total, or in only one sample, were discarded. Vaginal samples with fewer than 10,000 reads and/or features present in less than 20 samples were discarded. Four samples and all negative controls did not achieve sufficient sequence variants for downstream analysis leaving 238 samples that were included in the final analysis. Phylogenetic trees were constructed using MAFFT [39] for sequence alignment and FastTree [40] for tree construction. For the purposes of diversity analyses, samples were rarefied to a depth of 10,000 reads. Faith’s PD and the Shannon index were used to measure alpha diversity, and weighted Unifrac [41] was used to measure beta diversity.

### Lactobacilli DNA Extraction and qPCR Analysis

Known concentrations of *Lactobacillus iners* (ATCC 55195), *Lactobacillus crispatus* (ATCC 33820) *Lactobacillus gasser*i (ATCC 33323), and *Lactobacillus jensenii* (ATCC25258) genomic DNA extracted using isopropanol precipitation, was used to measure the detection limit of the species-specific 16S rRNA gene qPCR assays and to generate standard curves for quantifying assay results. DNA was previously extracted from vaginal swabs as described in [42] and along with lactobacilli genomic DNA was quantified using the Qubit 4 flurometer (Invitrogen Inc., Carlsbad, CA) with the Qubit dsDNA high-sensitivity assay (Invitrogen Inc., Carlsbad, CA). 10-fold serial dilutions (10^0^ to 10^8^ copies) were used to generate standard curves of extracted genomic DNA. The range of slopes for the qPCR assays was from −3.7 to −4.7, and r^2^ values were all > 0.99.

qPCR primers targeting the 16S rRNA genes of *L. iners, L. crispatus, L. gasser*i, and *L. jensenii* were previously developed in [43]. SYBR green-based qPCR assays were performed on a CFX96 Touch Real-Time PCR Detection System (Bio-Rad, Hercules, CA). The reaction mixture (20 µL) contained 10 µL iQ SYBR Green Supermix (Bio-Rad), 2 µL each of forward and reverse primers to a final primer concentration of 100 nM each except for the assay for *L. iners* which had used 4 µL each to a concentration of 200 nM., and 10 µL template DNA (10 ng). Temperature cycling for all assays was polymerase activation at 95°C for 3 min, followed by 40 cycles of amplification with denaturation at 95°C for 15 seconds, followed by annealing/ extension at 55 to 58°C for 1 min (*L. gasser*i, *L. iners* and *L. jensenii* at 55°C and *L. crispatus* at 58°C). Fluorescence was measured at the final step of each cycle. Following amplification, melting curve analysis was performed by heating at 0.5°C increments from 55 to 95°C, temperature was held for 5 s at each step, and fluorescence was acquired at each temperature increment. For each qPCR assay, vaginal swabs and extracted lactobacilli genomic DNA standards were run in triplicate, and the average values were used to calculate 16S rRNA gene copy number per 10 ng total vaginal swab DNA. Negative (no DNA water) controls were run with every assay to check for contamination.

### Virus-Like Particle Preparation, Library Construction and Sequencing

Virus-like particle (VLP) preparation was adapted from methods previously described in [44]. Briefly, vaginal specimens were resuspended in 200 μL of Digene transport media and an equal volume of SM buffer was added (50mM Tris-HCl, 8mM magnesium sulfate, 100mM sodium chloride, and 0.01% gelatin, pH 7.5, Fisher) and mixed by vortexing for 5 minutes. Specimens were centrifuged at 2000 x g and filtered using a 0.45 μM filter to remove intact cells and bacteria. Samples underwent lysozyme treatment (1ug/mL at 37C° for 30 minutes) (Sigma-Aldrich) to degrade remaining host cell and bacterial membranes. DNase digestion (Turbo DNase Buffer, Turbo DNase, Baseline Zero) (Ambion) was performed to remove contaminating bacterial and host DNA followed by heat inactivation of the DNase at 75C° for 15 minutes. Enriched VLPs were lysed with 10% SDS and 20mg/mL Proteinase K (Ambion) at 56C° for 20 minutes, followed by treatment with CTAB (10% Cetyltrimethylammonium bromide, 0.5 M NaCl, nuclease-free water, filtered through 0.22 μM filter) at 65C° for 20 minutes. Phenol: Chloroform: Isoamyl Alcohol (Invitrogen, pH 8.0) nucleic acid extraction was performed, the resulting aqueous fraction then washed with an equal volume of chloroform and concentrated through isopropanol precipitation. Extracted nucleic acid was aliquoted and stored at −80C° until use. NEBNext Ultra II FS DNA Library Prep Kit (New England Biolabs) was used for library construction with NEBNext Multiplex Oligos for Illumina Dual Index Primers (New England Biolabs). Following equimolar pooling, DNA libraries were sequenced on the Illumina NovaSeq platform (UR-GRC) generating an average of over 29 million 150bp paired-end reads per sample.

### Virome Analysis Pipeline

The VirusSeeker pipeline [45] was deployed on a Linux cluster on raw sequence data. Briefly, raw sequences from samples went through sample pre-processing steps that included adapter removal, stitching of reads, quality filtering, and CD-HIT was used to minimize sequence redundancy and define unique sequences (98% identity over 98% of the sequence length). Sequencing reads underwent human genome filtering, then unmapped reads were sequentially queried against a customized viral database comprised of all viral sequences in NCBI using BLASTn (e-value cutoff 1E−10), followed by BLASTx (e-value cutoff: 1E−3). False positive viral sequences were identified by successively querying the candidate viral reads against the NCBI NT database using MegaBLAST (e-value cutoff: 1E−10), BLASTn (e-value cutoff: 1E−10), and the NCBI NR database using BLASTx (e-value cutoff: 1E−3). All sequences that aligned to viruses were further classified into viral genera and species based on the NCBI taxonomic identity of the top hit.

### Statistical Analysis

Descriptive statistics were used to summarize the characteristics of the study population. Mean was used for continuous variables, and frequency or proportion was used for categorical variables. Alpha diversity was measured by Shannon’s diversity index and analyzed using a linear mixed effects model for bacteria data and a linear regression model for bacteriophage data [46]. Beta diversity was measured by the weighted UniFrac distance for bacteria data and by the Bray-Curtis distance for bacteriophage data [44, 47]. PERMANOVA was used to quantify dissimilarity in beta diversity [48]. Clustering was performed using partitioning around medoids, with the number of clusters estimated by maximum average silhouette width [49]. For differential abundance analysis, the relative abundance was first arcsine-transformed, and then univariate analysis was performed using mixed effects models for bacteria data and linear regression models for bacteriophage data. Univariate analysis of correlations between bacteria and bacteriophage was performed using Kendall’s rank correlation coefficient [50]. All analyses were performed in R and Prism version 8.3.0 for Windows (GraphPad Software, La Jolla California USA, www.graphpad.com) [51]. *P* values in the univariate analyses were adjusted for multiplicity using the Benjamini-Hochberg procedure [52].

## Results

### Cohort Characteristics

Fifty HIV-positive and 50 HIV-negative young, sexually active South African women ages 16-21 were recruited in Cape Town, South Africa between October 2012 and October 2014 as part of the University of Cape Town HPV-HIV study to investigate high-risk HPV persistence in HIV-positive women as previously reported [33, 42]. Vaginal swabs were self-collected at six-month intervals for up to six consecutive visits along with medical and sexual history, laboratory data and demographic information. Pap smears and STI testing were performed at the baseline visit and each year. HIV-positive subjects had an average CD4+ T cell count of 477.5 cells/μL, and 22 (44%) were not on ART at the baseline study visit. There was an increased risk of HPV for HIV-positive participants at baseline (OR 5.299, 95% CI 2.048 to 13.73). This cohort included 54 BV-positive and 46 BV-negative subjects at baseline (Table 1). The median age of both BV-positive and BV-negative women was 19. BV-positive subjects were more likely to have HPV (*p*=0.0940) and a higher prevalence of high-risk HPV subtypes (*p*=0.0005) compared to BV-negative subjects. BV-positive subjects also exhibited higher rates of abnormal pap smears (*p*=0.1181). These data demonstrate that our cohort behaves similarly to other published cohorts and was suitable for further study [53].

**Table 1:**
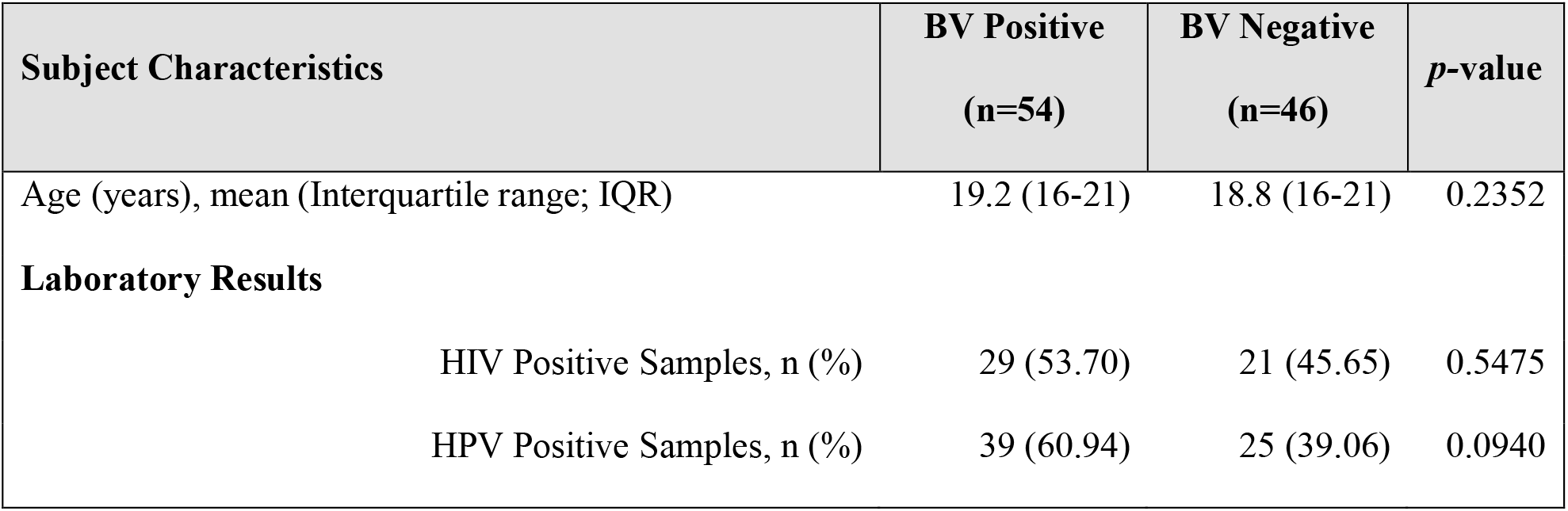
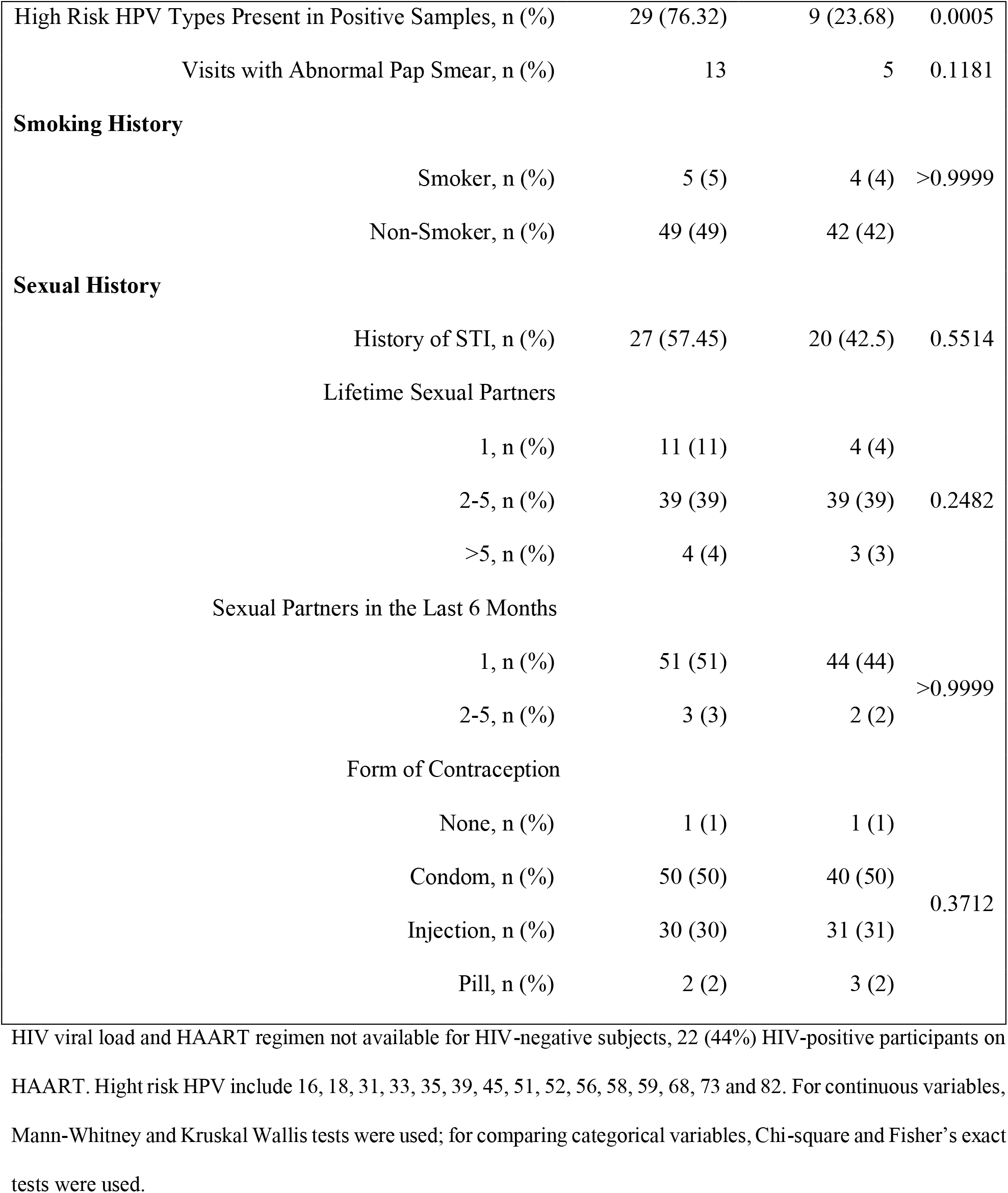
Cohort Characteristics.

### The FRT bacteriome clusters into community groups

Prior studies examining Western cohorts have identified five bacterial community state types (CSTs), of which type I-III and V were *Lactobacillus-*dominant while CST IV was comprised of polymicrobial communities [7]. However, studies examining the FRT bacteriome in African cohorts have revealed a different pattern in hierarchical clustering analysis, with more community groupings of high diversity bacteriomes, consistent with the increased prevalence of high diversity FRT bacteriomes in this population [4, 54]. To further asses the bacterial communities within the FRT bacteriome seen in African women, 253 vaginal swabs were processed and underwent 16S rRNA gene amplicon sequencing of the V3-V4 region [34]. Eleven samples did not amplify, and four failed to achieve sufficient reads for downstream analysis, leaving 238 samples for bacteriome analysis and sequence identification using QIIME2 (Fig.1A).

Hierarchical clustering analysis of all visits by community abundance and composition identified five unique bacterial community clusters named herein bacterial community groups (CG), which were distinguished by *Lactobacillus*-dominance (CG1 and 2) or higher diversity bacteriomes (CG3-5; Fig. 1A). The identified CG were compositionally different from conventional CSTs [7]. Similar to other African cohorts [7], the majority (n=173, 72.7%) of subjects had high diversity FRT bacteriomes with a low prevalence of *Lactobacillus*-dominant bacteriomes. Samples that were dominated by a single species, defined as >50% community composition, made up 42.0% of all samples and mostly clustered with CG1 and 2, while the remaining samples showed no individual dominant species and mainly clustered in CG3-5. CG1 (n=15, 6.3%), a low diversity FRT bacteriome, was comprised almost exclusively of *Lactobacillus* species (78%) that were unable to be further delineated by 16S rRNA gene amplicon sequencing (Fig. 1B). To further define the primary bacterial constituents of CG1, qPCR of 16S rRNA gene sequences from dominant FRT *Lactobacillus* species *L. iners, L. crispatus, L. gasseri* and *L. jensenii* [8, 54] was performed, revealing that approximately 70% of samples in GC1 were *L. crispatus*-dominant, similar to what has been previously described as CST I [55] (Fig. 1C). One vaginal swab in CG1 contained sufficient volume to only perform qPCR for *L. iners* and *L. crispatus*, and *L. crispatus* was most abundant (not shown). *L. jensenii* tended to predominate when present (Fig. 1C). CG2 (n=54, 22.7%) was *L. iners-*dominant, with a few samples showing notable amounts of *Gardnerella* and *Prevotella* as well (Fig. 1B). The second to largest and most diverse group, CG3 (n=64, 26.9%), consisted mainly of *Gardnerella, Prevotella* and *L. iners* (Fig. 1B). The smallest high diversity CG was CG4 (n=30, 12.6%), in which *Shuttleworthia* and *Gardnerella* were predominant. CG5 contained the largest number of samples (n=75, 31.5%) and was dominated by *Sneathia* and *Prevotella* (Fig. 1B). CG3, CG4 and CG5 exhibited significantly higher alpha diversity than CG1 and 2 (Fig. 2A; *p* = 0.0001, 0.0065, and <0.0001 respectively). Beta-diversity significantly differed between these five bacterial CG (*p*=0.0001; Fig. 2B).

**Figure 1:**
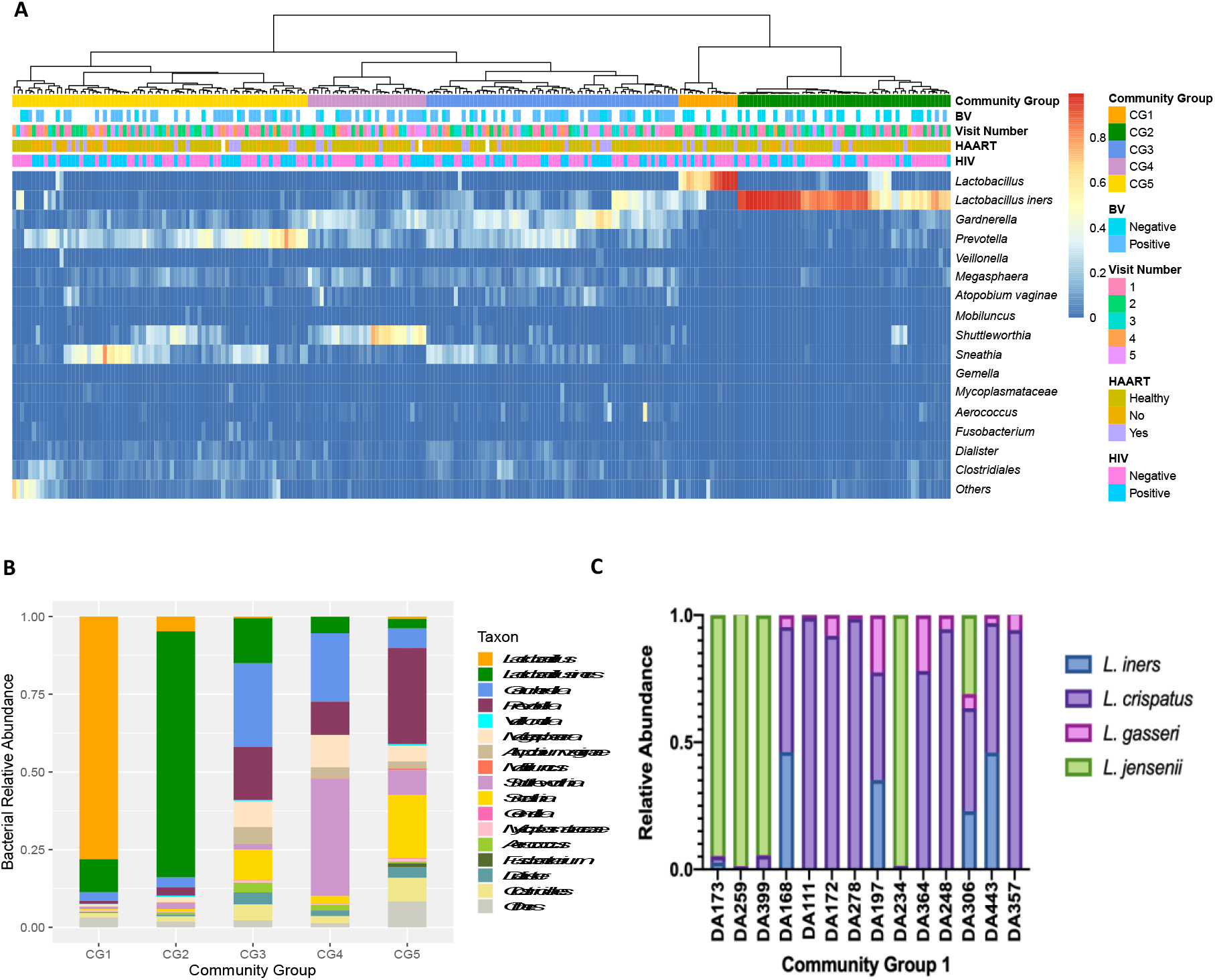
Bacteriome Profiling by Community Group of Self-Collected Vaginal Swabs from South African Women. (A) Relative abundance of 16 most frequent bacterial taxa (y-axis) by sample (x-axis), grouped by community group (CG), BV Status, visit number, HAART status and HIV status (color key shown). Percent abundance is indicated by gradient key. Using Ward’s linkage hierarchical clustering, samples clustered into five distinct bacterial community profiles called community groups (CG). (B) Average bacterial community group structure for each of the five community groups (x-axis) based on relative abundance (y-axis). (C) Bar plot showing the relative abundance of L. iners, L. crispatus, L. gasseri and L. jensenii bacterial species (relative abundance of 16S rRNA copies per 10 ng total DNA) as determined by qPCR of vaginal swabs that clustered into CG1.

**Figure 2:**
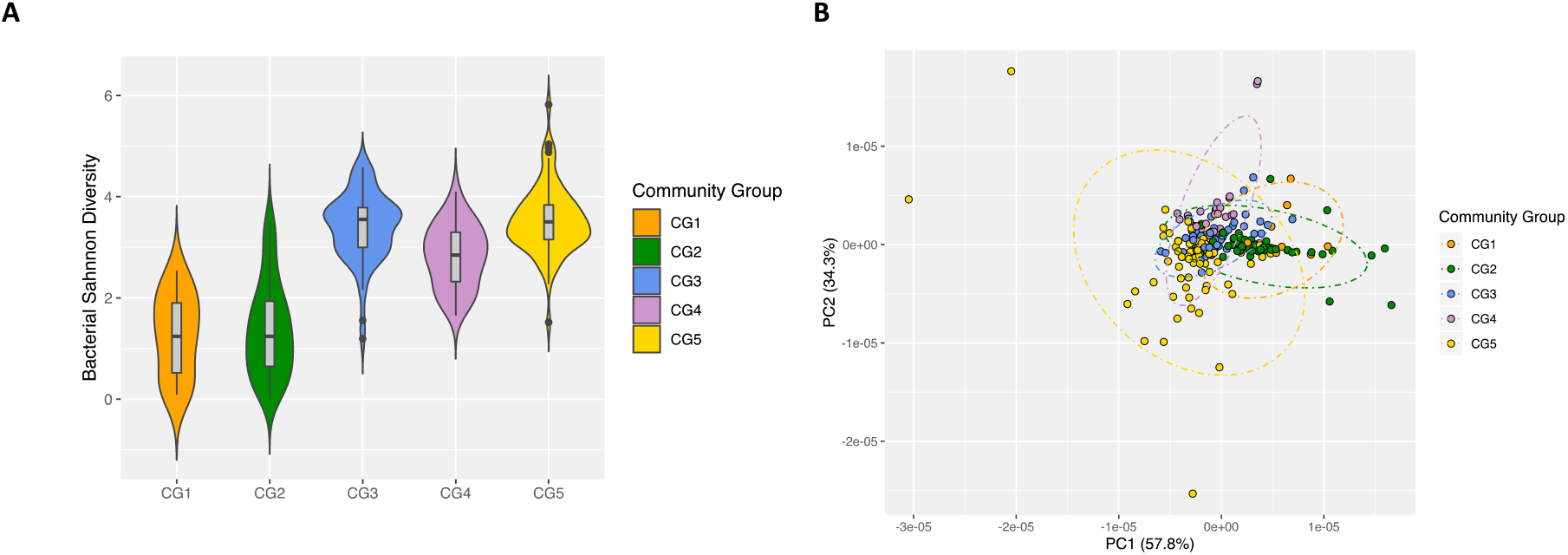
FRT Bacteriome Clusters into Distinct Community Groups That Differ by Alpha and Beta Diversity. (A) Bacterial Shannon diversity (y-axis) by CG (x-axis) as determined by linear mixed effects model. Center bar represents median; grey box bounded by upper/lower interquartile ranges (IQR); Whiskers represent range; Dots represent outliers; Color-filled areas are representative of density/distribution of diversity values. (B) Principle Coordinate Analysis (PCoA) plot of the weighted UniFrac distances colored by community group.

### Bacteriophages comprise the majority of the FRT DNA virome

While the FRT bacteriome has been well-studied, the FRT virome, especially bacteriophage populations, is relatively unknown. We therefore characterized the FRT DNA virome using a subset of baseline samples by enriching for virus-like particles (VLPs) from resuspended vaginal swabs and extracting viral nucleic acid [44]. Libraries were constructed and sequenced using the Illumina NovaSeq platform for 38 baseline samples, 14 of which were BV-negative and 24 were BV-positive. Resulting viral sequences underwent quality control, removal of bacterial and human reads, and then were assigned to known viral taxa using VirusSeeker, a BLAST-based NGS virome analysis pipeline [45].

On average there were 29 million reads per sample with 86.8% of them being high quality. The DNA eukaryotic virome was comprised almost entirely of *Papillomaviridae*. However, FRT bacteriophages were abundant. Samples contained sequences identified as belonging to *Myoviridae, Siphoviridae, Podoviridae, Inoviridae, Ackermannviridae, Microviridae, Lipothrixviridae, Plasmaviridae* and *Tectiviridae* bacteriophage families. Sequences assigned to members of the *Caudovirales* order, lytic tailed dsDNA bacteriophages, including *Myoviridae, Siphoviridae* and *Podoviridae*, were the most abundant in all samples regardless of BV, HAART, or HIV status.

Bacteriophage communities within the FRT clustered into two distinct, novel bacteriophage community groups based on composition and abundance that we have termed viral state types (VSTs) (Fig. 3A). VST1 represented 44.7% (n=17) of all samples while VST2 contained the remaining 55.3% (n=21). Bacteriophage Shannon diversity differed between VSTs (Fig. 3B), with VST2 having the highest diversity bacteriophage populations. The VSTs also grouped distinctly by beta diversity analysis (Fig. 3C, PERMANOVA, *p* =0.0001). Neither VST exhibited a dominant bacteriophage member. VST1 contained several samples with high relative abundance of *Rhodococcus viruses, Spounavirinae, phi29 virus* and *Biseptimavirus*-assigned bacteriophage reads. VST2 was comprised of a more even distribution. This is the first study to identify bacteriophage community groups in the FRT.

**Figure 3:**
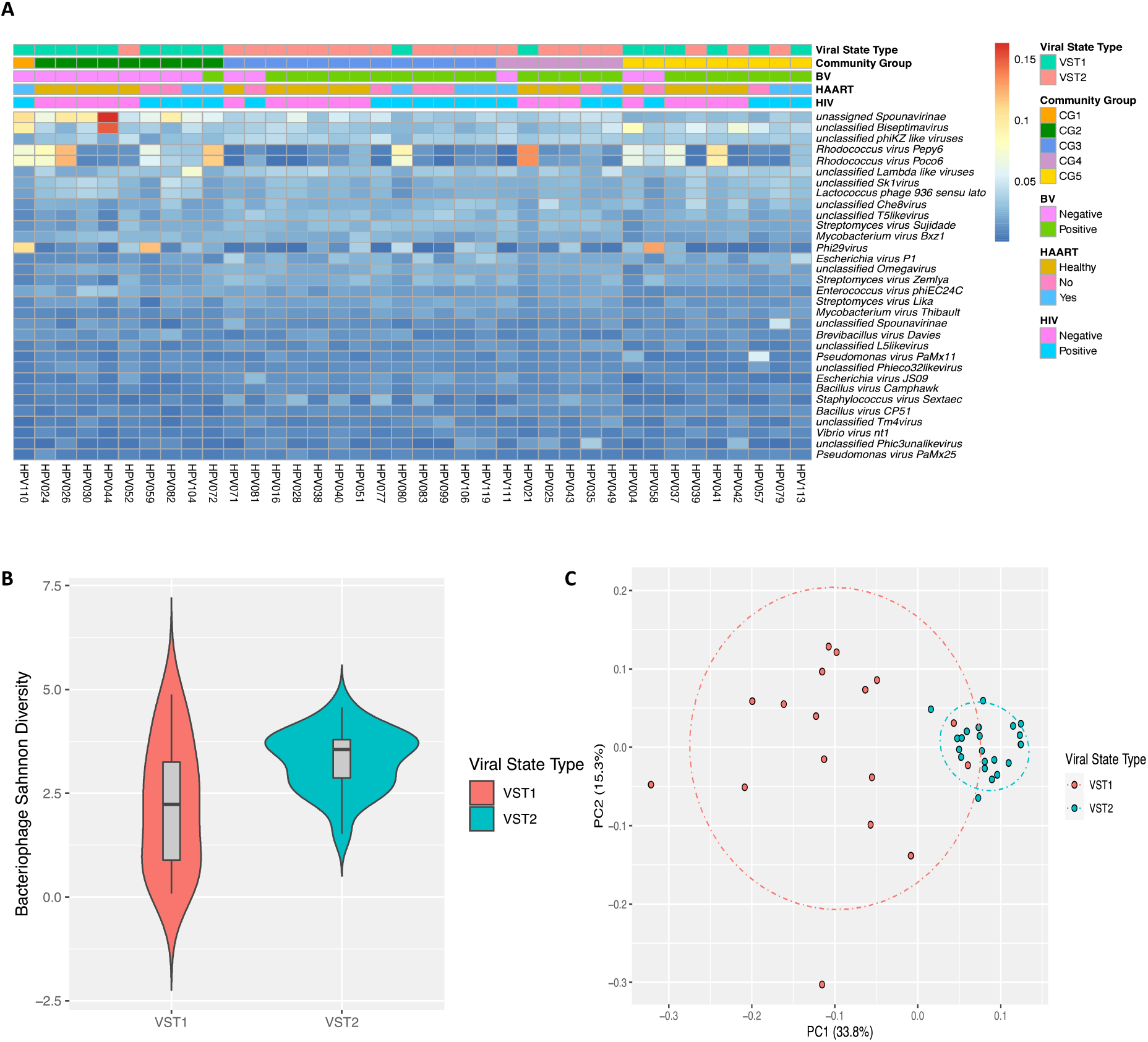
FRT DNA Bacteriophages Cluster into Two Unique Community Groups. Self-collected vaginal swabs were processed for DNA virome analysis by enriching for viral nucleic acid, libraries built, and sequenced. (A) Relative abundance of the 32 most frequent bacteriophage species (y-axis) by sample (x-axis). Ward’s linkage hierarchical clustering analysis was used to cluster samples into distinct bacteriophage community profiles called viral state types (VSTs). VST, BV Status, HAART status and HIV status (color key) are shown. Percent abundance is indicated by gradient key. (B) Bacteriophage Shannon diversity (y-axis) by VST (x-axis) as determined by linear regression model. Center bar represents median; grey box is bounded by upper/lower interquartile ranges (IQR); whiskers represent range; dots represent outliers; color-filled areas are representative of density/distribution of diversity values. (C) Principle Coordinate Analysis (PCoA) plots of beta diversity distances, as determined by Permutational multivariate analysis of variance, colored by VST.

### Transkingdom Associations within the FRT Microbiome of South African Women

Bacteriophage can directly impact bacterial composition and abundance through infection of their host. Therefore, we investigated the transkingdom associations between bacteriophage and bacteria in the FRT. The VSTs significantly correlated with bacterial CG (*p*=0.00015; Fig. 3A), with VST1 associated with CG1 and 2, the *Lactobacillus*-dominant groups, and VST2 associated with CG3 and 4, both higher diversity CG. CG5 contained samples belonging to both VSTs (Fig. 3A). These data indicate a strong association between bacteriophage communities and the host bacterial populations.

To further investigate specific bacteriophage-bacterial interactions, correlations between FRT bacterial composition and bacteriophage composition were identified by Kendell’s rank correlation coefficient. Reads assigned to bacteriophages *Bacillus virus Camphawk* and *Bacillus virus Pony*, which infect members of the *Bacillus* genus, positively correlated with the BV-associated bacteria *Gardnerella, A. vaginae, Prevotella, Sneathia* and *Dialister* (Fig. 4). Reads assigned to unclassified bacteriophages of the *E125* genus also positively associated with *Gardnerella* and *A. vaginae* (Fig. 4). A number of *Bacillus*-infecting phage including, *Bacillus virus Pony and Bacillus virus Staley*, were inversely associated with bacteria protective from BV, particularly *L. iners* (Fig. 4). Interestingly, in this cohort, bacteriophage associations revealed *Veillonella* more closely grouped with *Lactobacillus* rather than BV-associated bacteria despite the known role of *Veillonella* in lactose fermentation and higher diversity FRT microbiomes [7]. These data suggest that bacteriophages directly or indirectly interact with FRT bacterial populations in disease states, and identify putative FRT bacteriophage-host networks that may play a role in development and maintenance of BV.

**Figure 4:**
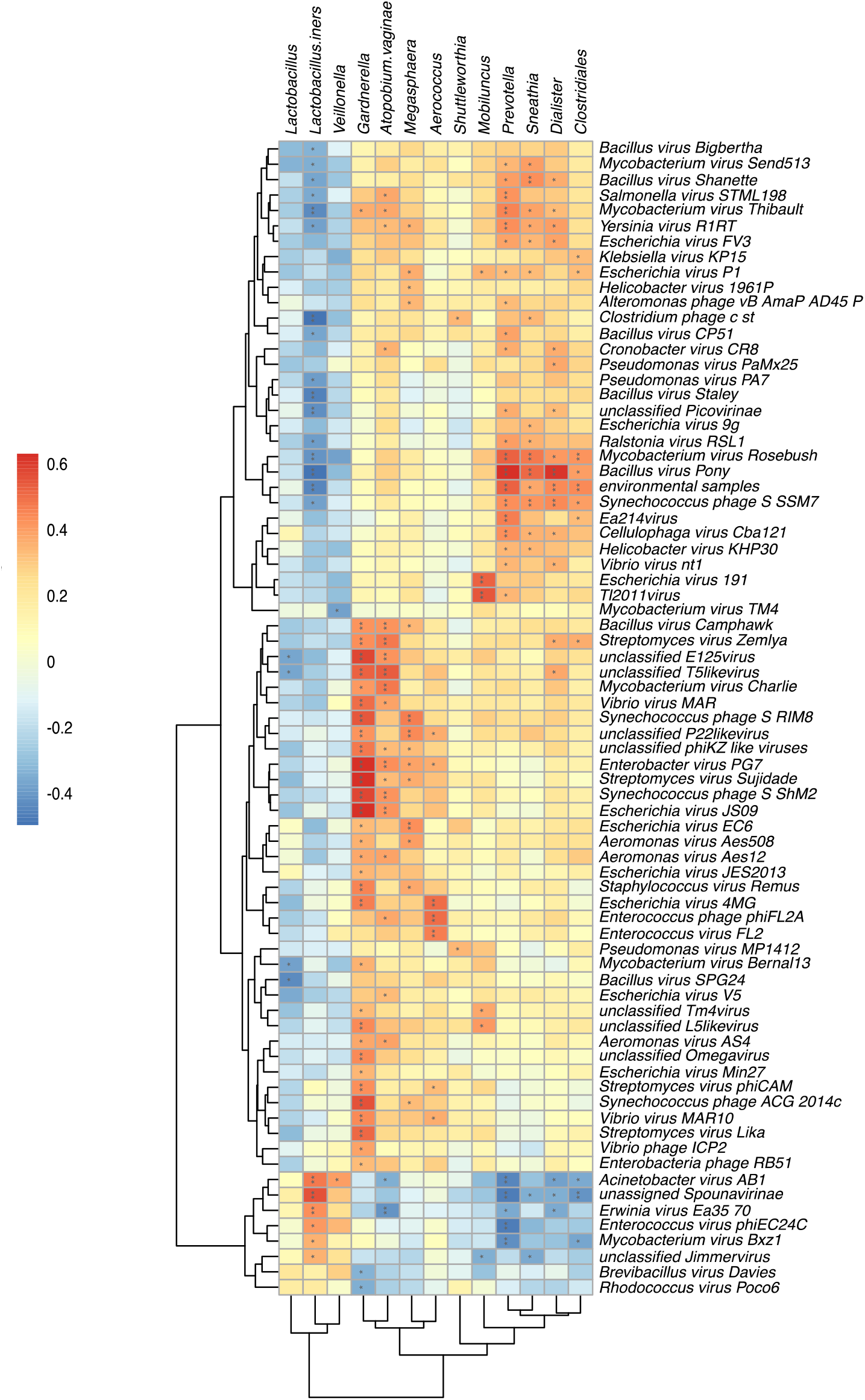
Transkingdom Associations within the FRT of South African Women. Heatmap of estimated Kendall’s correlation coefficients between FRT bacterial taxa (x-axis) and sequences assigned to bacteriophage (y-axis). Multiple comparisons correction by the Benjamini-Hochberg procedure. * indicates p<0.05, ** p< 0.01. Magnitude and sign of the Kendall’s rank correlation coefficient is indicated by gradient key. Red indicates positive correlations; blue indicates negative correlations.

### Effects of Bacterial Vaginosis on the FRT Virome

We next examined the impact of different disease states on this cohort. BV is a clinically significant condition with high morbidity characterized by high bacterial diversity [12, 13]. Our data showed significant associations between high diversity bacterial CG and VSTs, and further suggested specific bacteriophage interactions with BV-associated bacteria. Similar to published cohorts [2], BV in our cohort was positively associated with increased bacterial alpha diversity compared to healthy subjects (*p*=0.0001). The high diversity CG3-5 (*p*=0.0001) also correlated positively with BV. Bacterial taxa including *Gardnerella, Prevotella, Sneathia and Megasphaera* (*p*=0.000439; Fig. 5A) were linked to clinical BV status, corroborating distinct bacterial signatures associated with BV. Of particular clinical relevance, we additionally sought to identify specific bacterial taxa that were predictive of initiation and recovery from BV. Relative abundance of bacterial genera less frequently observed in clinical BV, Aerococcus (p=0.00216) and Gemella (p=0.00746), was significantly increased among participants who recovered from or transitioned to BV, suggesting members of this genera as possible regulators of FRT bacteriome structure in health and BV.

**Figure 5:**
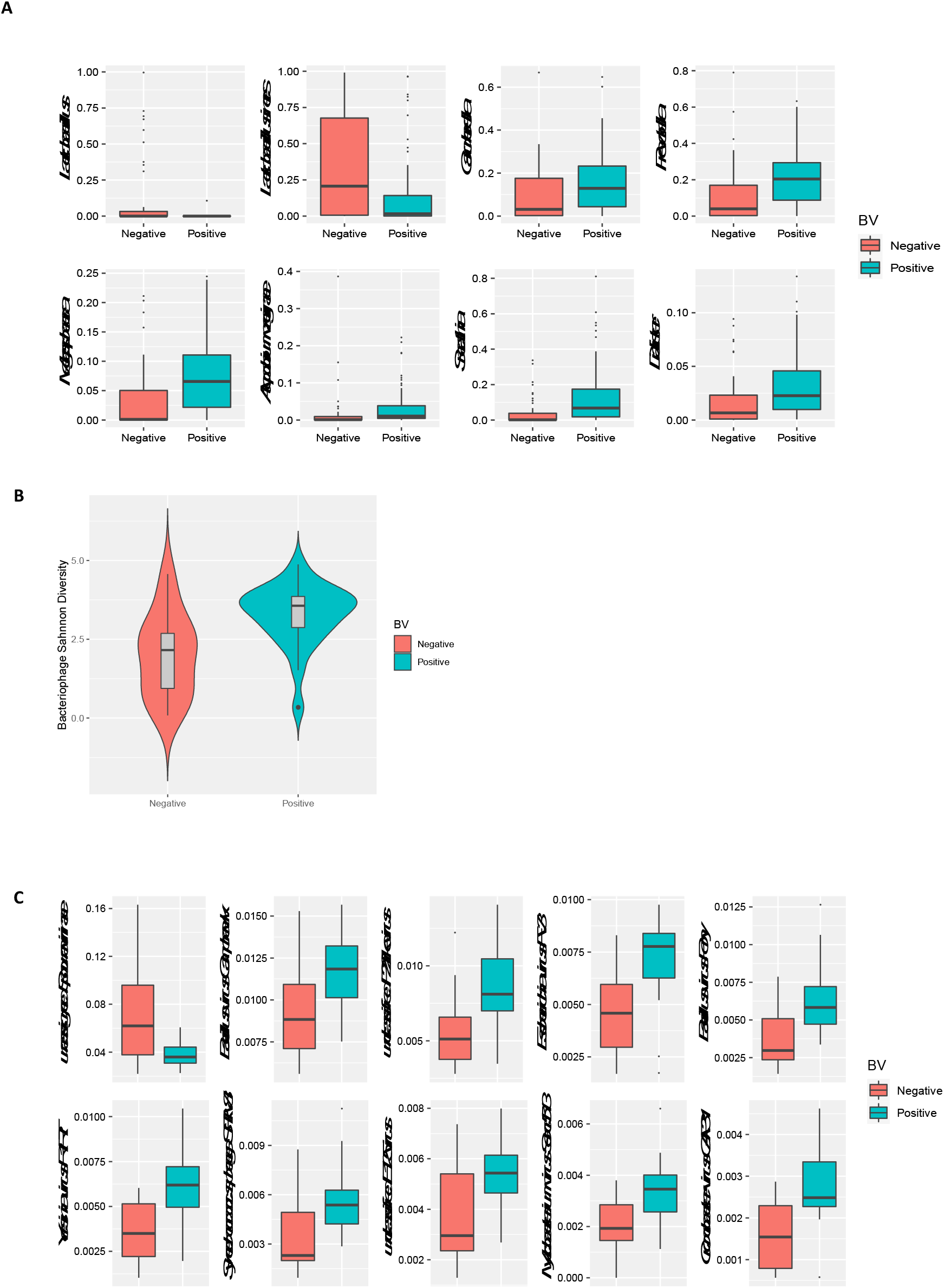
Discriminant FRT Bacterial and Bacteriophage Species Associated with Bacterial Vaginosis. (A) Discriminant bacterial taxa by BV status was determined by univariate analysis using mixed effects models. Relative abundance is represented on the y-axis, BV-negative subjects are shown in orange and BV-positive in green (x-axis). (B) Bacteriophage Shannon diversity (y-axis) by BV status (x-axis) as determined by a linear regression model. (C) Discriminant bacteriophage species by clinical BV diagnosis was determined by univariate analysis using a linear regression model. Relative abundance is represented on the y-axis, BV-negative subjects are shown in green and BV-positive in orange (x-axis).

Since clinical BV was associated with significant changes in FRT bacterial composition and correlated with bacterial CG, and bacteriophage VSTs also correlated with bacterial CG, we sought correlations between bacteriophage populations and clinical BV. Regression analysis accounting for HIV status and VST revealed bacteriophage Shannon diversity significantly differed by clinical BV diagnosis (*p*=0.0193) (Fig. 5B). We then identified specific bacteriophage taxa differentially abundant in the FRT of BV-positive and -negative women. *Bacillus*-infecting bacteriophages are known to belong to the *Herelleviridae* and *Podoviridae* families [56, 57]. Members of these families, including *Bacillus virus Camphawk* and *Bacillus virus Pony* (Fig. 5C), which were significantly associated with BV diagnosis (*p*= 0.019058 and *p=*0.014546, respectively). BV diagnosis also strongly correlated with the bacteriophages *Escherichia virus FV3* and *unclassified E125 virus*, which are known to infect the BV-associated bacteria *Escherichia coli* and *Burkholderia*, respectively [58, 59]. Together, these data uncover a link between highly diverse FRT bacteriophage populations, a distinct subset of bacterial hosts, and BV.

### The Effect of HIV and HPV on the FRT Microbiome

We also examined the effect of HIV on the FRT microbiome in this cohort. We found no significant difference in bacterial richness, alpha or beta diversity between HIV-positive and HIV-negative subjects. Further, there were no significant alterations in bacteriome or bacteriophage diversity by HIV status. Thus, FRT bacterial and bacteriophage communities were not detectably altered by HIV infection in this cohort, suggesting that localized infections may play a more important role in the FRT composition than systemic infections.

We also examined the relationship between HPV, the main eukaryotic virus found, and bacterial populations. Upon examination of 63 HPV-positive and 37 HPV-negative baseline samples, there were no significant associations between HPV infection and bacteriophage or bacterial diversity. Analysis of HPV subtypes revealed increased bacterial alpha diversity in HPV6-positive subjects (*p*=0.0181), suggesting certain HPV subtypes may directly or indirectly benefit from the presence of higher diversity bacterial populations, although this analysis may have been underpowered for less prevalent subtypes.

## Discussion

The FRT is a dynamic ecosystem in which bacteriophage and bacterial communities establish complex connections that influence the host environment and the physical manifestation of gynecological diseases. Bacterial population shifts are well-established contributors to FRT disease states including BV [7]. However, whether shifts in the FRT viral community occur in disease states concurrently with bacterial community perturbation was unknown. This study is the first to offer a comprehensive characterization of both the FRT bacteriome and DNA virome, with a particular emphasis on bacteriophage composition and transkingdom interplay, utilizing a South African cohort of BV-affected women.

Globally, the most common clinical presentation of vaginal bacterial dysbiosis is BV, a condition which poses a significant threat to female reproductive health and STI acquisition [2, 12, 13]. Here, similar to other studies, we find that women suffering from BV have distinct and compositionally diverse FRT bacteriomes defined by a loss of *Lactobacillus* dominance and gain of facultative anaerobes [7]. However, for the first time, our data revealed novel bacteriophage communities we have termed VSTs, which correlated to bacterial communities and clinical diagnosis of BV. Although bacteriophages have been studied at other mucosal sites in humans including the gut [60], this is the first description of bacteriophage community groupings and may be unique to the FRT environment due to the distinctive bacterial communities present. Few prior studies have examined the bacteriophage populations in the FRT. Nonetheless, in contrast to our findings, one group examining FRT bacteriophage populations found no distinct bacteriophage community structures within their cohort, nor differences in bacteriophage composition among the bacterial communities [4]. Similar to our study, they also utilized a South African cohort; however, they focused analysis on the *Caudovirales* family rather than all bacteriophage populations, which likely impacts differences observed. Additionally, our findings contrast with those preciously described by Jakobsen et. al [23], who found no significant difference in viral nor bacterial alpha diversity between BV-positive and BV-negative samples in a cohort of Danish women undergoing IVF treatment for non-female factor infertility. However, their analysis was limited by the cross-sectional nature of the study and meager modest sequencing depth. Our viral sequencing depth of 29 million reads per sample was greater than 640-fold higher than that employed in Jakobsen et al. (average of 44,686 reads per sample) and is likely a major contributor to our unique findings. Additionally, Zhang et al. briefly characterized the vaginal microbiome in pregnant women finding absence of Microviridae and Herelleviridae in a pregnant woman without vaginitis [24]. This study was limited in statistical analysis due to the pooling of subjects into size sequencing libraries and similar to Jakobsen et al., Zhang et al. used a sequencing depth an order of magnitude lower that what was employed in our current investigation. Validation of our novel FRT VSTs using other cohorts is currently underway.

As the etiology of rapid microbiome shifts between health and BV disease states remains unclear, we sought to identify specific bacterial taxa associated with recovery and transition to BV that could be involved in these dynamic changes. We observed a significant increase in relative abundance of the gram-positive facultatively anaerobic cocci *Aerococcus* and *Gemella* during transition to and from clinical BV. However, previous literature has recognized these bacterial genera as being less frequently detected during clinical BV [61]. This could indicate these taxa as important facilitators of disease-associated microbiome composition and as clinical biomarkers of BV. Similar to what is seen in the gut, these bacteria may be assuming the role of “primary species” commonly seen after an environmental disturbance, whereby fast-growing facultative anaerobes transiently bloom and are essential for the presence of other taxa along with ecological diversity and structure [62]. Interestingly, *Enterococcus virus FL2* A and *Enterococcus phage phiFL2A* both were found to be significantly associated with the transitory bacteria *Aerococcus*. Because of the sequence similarity between *Aerococcus* and *Enterococcus* bacteria, along with the identification of these bacteriophage prior to the introduction of more sensitive next generation sequencing technologies, it is probable that *Enterococcus virus FL2* A and *Enterococcus phage phiFL2A* have a misidentified host [63]. These bacteriophages may actually target *Aerococcus* species, facilitating their significant role in recovery and transition to BV. A larger longitudinal patient population would be needed to further distinguish between specific taxa associated solely with BV incidence vs BV recovery and the roles they play in these FRT microbiome transitions.

Our analysis also is the first to identify discriminant bacteriophage taxa by BV status and assess transkingdom associations in the FRT. Correlation analysis between bacterial taxa and assigned bacteriophage species identified bacteriophages that positively correlated with BV-associated bacteria and inversely correlated with *Lactobacillus*, suggesting that transkingdom interactions between bacteriophages and bacterial species could be the driver of BV-associated bacterial community alterations. *Bacillus virus Camphawk* and *Bacillus virus Pony*, previously demonstrated to be lytic to *Bacillus* members [64, 65], were associated with both BV and BV-associated bacteria. In the gut, *Bacillus* strains have been shown to antagonize enteropathogenic bacteria, while concurrently promoting the growth of *Lactobacillus* [66]. If *Bacillus* species act in a similar manner in the FRT, the lytic nature of the *Bacillus* bacteriophages *Bacillus virus Camphawk* and *Bacillus virus Pony* could at least partially explain the shift in vaginal microbiota away from *Lactobacillus* species and toward more diverse bacterial species, including the facultative anaerobes seen in BV. *E. coli* and *Burkholderia* bacteriophages *Escherichia virus FV3* and *unclassified E125* virus, respectively, were also associated with BV. While less predominant than other bacterial species, *E. coli* and *Burkholderia* are also implicated in BV-associated bacterial communities [1, 2, 58]. *Escherichia virus FV3* and *unclassified E125* virus may act to regulate bacterial abundance and community composition in BV via a predator-prey relationship to allow for growth of primary BV-associated bacterial members such as *Gardnerella* and *Prevotella* [22, 67]. BV risk factors such as new or multiple sexual partners provide a plausible mechanism for introduction of novel bacteriophage that could target and deplete commensal FRT bacteria [68, 69].

Interestingly, we did not find any significant association between sequences assigned to bacteriophage know to infect the more common BV-associated bacteria, including *Gardnerella* or *Prevotella*, by either BV-discriminant taxa or transkingdom analysis. The absence of bacteriophage that infect hallmark BV bacteria such as *Gardnerella vaginalis* in our analysis may be attributable to the high proportion of *Gardnerella* species that contain CRISPR/Cas-9 bacteriophage defense loci, making them more resistant to bacteriophage infection and establishing a uneven bacteriophage burden between bacteria in BV [70]. These data suggest a very active, dynamic environment of bacteriophage warfare against numerous bacterial hosts to regulate the bacterial populations. Further *in vitro* studies will be necessary to determine host range and bacteriophage lifestyle. A thorough examination of the FRT microbiome in the context of BV contributes to a deeper understanding of BV pathogenesis.

While we focused on BV as a common FRT-localized disease, we also were interested in investigating the effect of other disease states on the FRT microbiome. In addition to BV, we studied the impact of HIV infection on the FRT but observed no difference in FRT bacterial or bacteriophage diversity based on HIV status. One possible explanation is that HIV, as a systemic disease, does not have a major impact on the localized FRT mucosal environment. Prior literature shows that acute HIV infection leads to distinct changes in local inflammatory marker profiles, including elevation of pro-inflammatory cytokines IL-1α, IL-1β, and IL-6 [4, 54] however, it remains unclear if this initial inflammatory response persists long-term. Alternatively, we may not have detected differences due to the well-controlled nature of HIV infection in this cohort. We previously showed that immunocompromise was a major factor affecting enteric microbiome diversity in HIV infection [44], a finding replicated at other mucosal sites [71], suggesting increased likelihood of observing microbiome alterations with immunodeficiency. Since the mean CD4+ T cell count for this cohort was 478 cells/μL, and no subjects were known to be immunosuppressed, this could explain the dearth of associations of the FRT microbiome with HIV seen in our study.

Limitations to this study include the cross-sectional nature of the virome analysis, preventing speculation on the longitudinal impact of bacteriophage changes, and low RNA integrity of the samples, blocking assessment of the FRT RNA virome. The initial limited patient consent also precluded significant *in vitro* validation of bacteriophage-bacterial pairs. Finally, there may be inaccuracies or biases in sequence assignments.

## Conclusions

In this retrospective longitudinal study, we performed a novel in-depth investigation of the FRT virome and bacteriome using a cohort of young, sexually active, South African women. We discovered significant alterations in FRT bacterial and bacteriophage diversity and community structure associated with BV. Transkingdom analysis revealed associations of specific bacteriophages with bacteria protective of and associated with BV. This study is the first to describe VST structure within the FRT and its associations with bacterial diversity and composition. The nature of the FRT in health and disease is both complex and dynamic and our findings provide insight into putative interactions between bacteriophage and bacteria that may contribute to development and maintenance of FRT dysbiosis. Further studies are needed to investigate direct mechanisms employed by bacteriophages to promote dysbiosis.

## List of abbreviations

BV: Bacterial Vaginosis
CG: Bacterial Community Group
HIV: Human Immunodeficiency Virus
FRT: Female Reproductive Tract
NGS: Next Generation Sequencing
STI: Sexually Transmitted Infection
VST: Viral State Type

## Ethics approval and consent to participate

This study was reviewed by the Research Subjects Review Board of the University of Rochester and granted human exemption status.

## Informed Consent

Informed consent was obtained from all subjects involved in the study

## Authors’ contributions

Conceptualization and Methodology: CLM and FSM; Formal Analysis – MS and AG; Investigation: FSM, BB, AW; Resources and Sample Acquisition: TM, ALW, LGB, DHA; Data curation: FSM, BB, AW; Writing – original draft: FSM; Writing – reviewing and editing: all authors; Visualization: FSM, MS, AG; Supervision: CLM; Funding acquisition: CLM.

## Acknowledgments

We thank the study subjects for their participation as well as study team nurses and personnel. We also would like to thank James Java, Ph.D. for assistance with the VirusSeeker virome pipeline and Cassandra Newkirk for experimental support. We thank the Center for Integrated Research Computing (CIRC) at the University of Rochester for providing computational resources and technical support. Microbiome and virome sequencing in this study was completed by the University of Rochester Genomics Research Center (GRC).

